# A pleiotropic *EPAS1* enhancer mediating Tibetan adaptation to hypoxia is active in adipocytes

**DOI:** 10.64898/2026.06.11.730447

**Authors:** Alexis G. Thornburg, Soo-Young Park, Li Zhang, Ivy Aneas, Débora R. Sobreira, Isabella M. Salamone, Noboru J. Sakabe, Kathryn M. Farris, Zachary T. Weber, Olivia A. Gray, Jennifer Yoo, Hae Kyung Im, Anna Di Rienzo, Marcelo A. Nóbrega

**Author notes:** For inquiries, please contact Alexis G. Thornburg, Anna Di Rienzo, or Marcelo A. Nóbrega.

## Abstract

In response to hypoxic stress at high altitudes, variation at the *EPAS1* locus has experienced strong selection in Tibetans. Functional dissection of the selection signals at this locus identified ENH5, an enhancer within the adaptive haplotype that has a blunted response to hypoxic stress in Tibetans. ENH5 was shown to be pleiotropic in several tissues related to hypoxia response, suggesting that a possible mechanism behind the strong selection signatures could be adaptive pleiotropy. Tibetans not only experience hypoxic conditions, but also cold temperatures due to the altitude and climate of the Tibetan Plateau. However, it is unclear whether cold temperatures affect ENH5 activity possibly contributing to the selective pressure at this locus. Here, we further characterized the role of ENH5 in subcutaneous white adipose tissue, an important tissue type that regulates body temperature in response to cold temperatures by releasing stored fat as heat through a process called thermogenesis. In this work, we investigated the role of ENH5 in adipocytes using ENH5 knockout mice (ENH5 KO), which phenocopy the reduced activity of the Tibetan allele. We show that ENH5 KO mice at normoxia and room temperature do not have significant differences in organismal phenotypes related to adiposity and metabolism compared to WT mice on a high fat diet. However, we detected effects of ENH5 conditional on thermogenic stimulation and hypoxia exposure, independently, in adipocytes cultured *in vitro*. Under either of these conditions, ENH5 KO has stronger differential expression of key genes involved in thermogenesis activity and adipocyte differentiation compared to WT. This differential response to thermogenic stimulation expands on the pleiotropic effects of the Tibetan ENH5 allele(s), in addition to those previously shown in well-established hypoxia-responsive tissues. Our results raise the possibility that pleiotropic effects of ENH5 may implicate unforeseen mechanisms, such as cellular energetics and thermogenesis, possibly contributing to the phenotypic adaptation to high altitude in Tibetans.

## INTRODUCTION

Human populations have evolved diverse genetic adaptations in response to varied environmental pressures encountered across the globe. Key adaptations include physiological, metabolic, and immunological changes driven by natural selection acting on genetic variants^1^.

Adaptations to life in high altitude is a particularly well characterized example. The human populations with a long-term history of high altitude residence (e.g., Tibetans, Andeans, Ethiopians) exhibit distinct genetic modifications affecting hemoglobin concentration and hypoxia pathways, vascular biology and mitochondrial function^2^. Interestingly, these distinct human populations seem to rely on genetic adaptations at different genetic loci favoring their life in high altitude. Variants in *CBARA1, VAV3, ARNT2,* and *THRB* underlie adaptations in Ethiopians living in high altitude^2^, while Andean populations show signatures of selection in the locus encoding PHD2, which is part of the HIF pathway, and *PRKAA1*, which plays a role in cellular energy balance and metabolism^3^. Genetically, Tibetans exhibit strong natural selection in genes like *EPAS1* and *EGLN1*. The selection signal observed at the *EPAS1* locus remains one of the strongest, most reproducible examples of genetic adaptation in humans^4–7^, and has been the focus of several follow up adaptation studies^7–10^.

Tibetans inherited unique genetic variants in the *EPAS1* locus, notably through ancient interbreeding with Denisovans, an archaic human species^11^. The variants under selection at this locus are associated with lower hemoglobin (Hb) concentration in Tibetans^4,5,7^. This lower Hb phenotype has been associated with better reproduction outcomes in Tibetan women^12^, suggesting that these selected variants may act on Tibetan phenotypes to provide a significant survival advantage for living in the Tibetan Plateau. Some other unique phenotypes exhibited by Tibetans are enhanced oxygen utilization^13^, low pulmonary artery pressure^14,15^ and lower rates of pulmonary hypertension^14^. However, elucidating the full suite of phenotypes associated with selection signals and dissecting the specific phenotypes that confer a selection advantage remains an open area of research. While advances in human population genomics have facilitated the identification of beneficial alleles by means of genome scans for selection signatures^16–18^, functional and phenotypic dissection of these selection signatures is needed to provide insights as to how natural selection shaped human physiology at high altitude.

Contrasting with the locus heterogeneity seen in different human populations that rely on different genes for their adaptations to life in high altitude, several mammalian species that live in the Tibetan Plateau also seem to rely on coding and noncoding genetic variants within the *EPAS1* locus, as do humans. These species include cattle^19^, dogs^19,20^, sheep^19,21^, horses^19,22^, and pikas^23^. This unusual architecture raises the possibility that aspects of the Tibetan Plateau environment limit the scope of genes that are involved in adaptations to high altitude, compared to other high-altitude regions across the globe. Our hypothesis is that the transcription factor encoded by *EPAS1*, i.e. HIF-2α, has functions beyond oxygen sensing and may also have been adaptive to life in Tibet. Among unique environmental differences between Tibet and other high-altitude regions, the Tibetan Plateau exhibits colder temperatures. For example, the average temperature in winter is approximately -9°C in the Tibetan Plateau^24^, while the average temperature in winter is approximately 10°C in the Andean Plateau^25^ and 8°C in the Ethiopian highlands^26^. Therefore, more extreme cold temperatures in the Tibetan Plateau could be a selective pressure acting on Tibetans in addition to hypobaric hypoxia.

Recent work investigating the genetic and molecular mechanisms underlying human Tibetan adaptation to high-altitude hypoxia identified a pleiotropic, hypoxia-sensitive enhancer (ENH5) within *EPAS1* that is disrupted by Tibetans’ adaptive alleles. High-altitude alleles reduce the activity of ENH5 across several tissues and cell types, leading to decreased *EPAS1* expression and attenuated transcriptional responses to acute and sustained hypoxia. Functional analyses in mice, who share the evolutionarily conserved *Epas1* ENH5, reveal that loss of ENH5 results in broad dysregulation of *Epas1* targets in heart, lung, kidney, and adrenal gland. These data suggest that the strong signature of selection at *EPAS1* in Tibetans is explained, at least partially, by the pleiotropic effects conferred by enhancer disruption, affecting multiple genes across multiple tissues^9^.

This prompted us to explore if the Tibetan ENH5 haplotype has an effect in response to cold temperature. We focused on assessing the role of ENH5 in adipocytes, an important cell type for regulating body temperature in response to cold temperatures. Adipocytes can produce heat through non-shivering thermogenesis, where β_3_ adrenergic receptor (β_3_-AR) activation leads to aerobic respiration being uncoupled from ATP production to release energy as heat^27–29^. While brown adipose is the primary fat depot for thermogenic response, it is primarily present in infants to regulate body temperature and is substantially reduced into adulthood^30,31^. However, subcutaneous white adipose can undergo a process called browning, where this fat depot increases their thermogenic capacity in response to β_3_-AR activation^27–29^. Interestingly, subcutaneous white adipocytes in humans have also been shown to increase their thermogenic response to cold temperatures during seasonal changes^32^, suggesting that this fat depot is involved in adaptation to environment. Additionally, HIF-2α has been shown to regulate thermogenesis in subcutaneous white adipocytes, where HIF-2α activation in response to oxygen consumption during cold stress acts as a negative regulator to suppress sustained thermogenic activity^33^. This led us to speculate that the Tibetan ENH5 haplotype might be affecting response to cold stress in subcutaneous white adipocytes due to this fat depot’s dynamic ability to respond to cold stress. In light of HIF-2α having been shown to regulate thermogenesis and since the Tibetan ENH5 has blunted enhancer activity, we would postulate that the Tibetan ENH5 would blunt the effect of HIF-2α as a negative regulator of thermogenesis, thus maintaining thermogenic activity in response to cold stress. This response could perhaps allow Tibetans to maintain body temperature in response to cold stress. However, maintained thermogenic activity can lead to a depletion of energy reserves^34^, and could thus be a maladaptive phenotype particularly in hypoxic conditions. On the other hand, blunted thermogenic activity would reduce heat production during cold exposure, allowing conservation of energy and minimizing metabolic demands^35^, which could be an adaptive phenotype. For populations in cold environments, this adaptation helps preserve body fat and calories and supports survival when food resources are scarce^36^. Therefore, regulating thermogenic activity to balance heat production and energy expenditure could be a phenotype influenced by this selected locus in Tibetans. Using an ENH5 knockout (ENH5 KO) mouse to model the Tibetan ENH5 haplotype, we set out to understand if and how thermogenic regulation in subcutaneous white adipocytes (hereafter referred to as adipocytes) may be influenced by this selected locus.

In our work, we first established that ENH5 has activity in adipocytes, providing further evidence of the pleiotropic role of this enhancer. Using ENH5 KO mice, we found ENH5 does not impact fat composition or metabolism *in vivo*, but it does have conditional effects in adipocytes cultured *in vitro* in response to thermogenic stimulation and hypoxia independently. Under each condition, ENH5 KO has stronger differential expression of genes involved in aerobic respiration and adipogenesis compared to WT, suggesting less energy is released as heat and fewer adipocytes are generated to respond to thermogenic stimulation. Therefore, ENH5 KO results in lower thermogenic activity, suggesting that the Tibetan ENH5 haplotype favors energy conservation over energy expenditure under cold temperatures.

## RESULTS

### ENH5 is active in adipose tissue and is conserved in sequence and function between humans and mice

The strong selection signal at the *EPAS1* locus, particularly at ENH5, observed in Tibetans who experience cold as well as hypoxic stress led us to explore a pleiotropic role of this enhancer. One tissue that is important to cold temperature response is adipose tissue, which can break down stored fats and release this energy as heat. To explore a role of ENH5 in thermogenesis, we looked at the chromatin landscape of this locus in adipose tissue. ENH5 is found to be in open chromatin, similarly to lung and kidney tissue, two tissue types where the activity of ENH5 has previously been established^9^ (Figure 1A). Knowing that ENH5 is in open chromatin in adipose tissue, we assessed if ENH5 is an active enhancer in this tissue type. Using 3T3-L1 cells, we tested the activity of the high-altitude and low-altitude alleles of ENH5 and found that both alleles are active under normoxia and hypoxia (1% O_2_, 48 hours). Additionally, we found that ENH5 has allele-specific responses under normoxia and hypoxia, where the high-altitude allele has blunted activity compared to the low-altitude allele in both conditions (Figure 1B). These allele-specific results match previous data showing blunted activity of the ENH5 high-altitude allele relative to the low-altitude allele in several tissue types^9^. To further explore the possible pleiotropic role of ENH5 in adipose tissue, we used a previously established ENH5 KO mouse line^9^ which models the reduced activity of the ENH5 high-altitude allele. This ENH5 KO mouse is an adequate model for investigating the pleotropic role of ENH5 since ENH5 is conserved at the sequence level between humans and mice (Figure 1C). Additionally, the mouse ENH5 is an active enhancer in 3T3-L1 cells under normoxic and hypoxic conditions (1% O_2_, 48 hours), thus indicating conservation of function between mouse and human ENH5 (Figure 1D). This mouse model harbors a homozygous 1kb deletion of ENH5, allowing us to interrogate the functional role of ENH5 *in vivo* and *in vitro* (Figure 1E).

**Figure 1.**
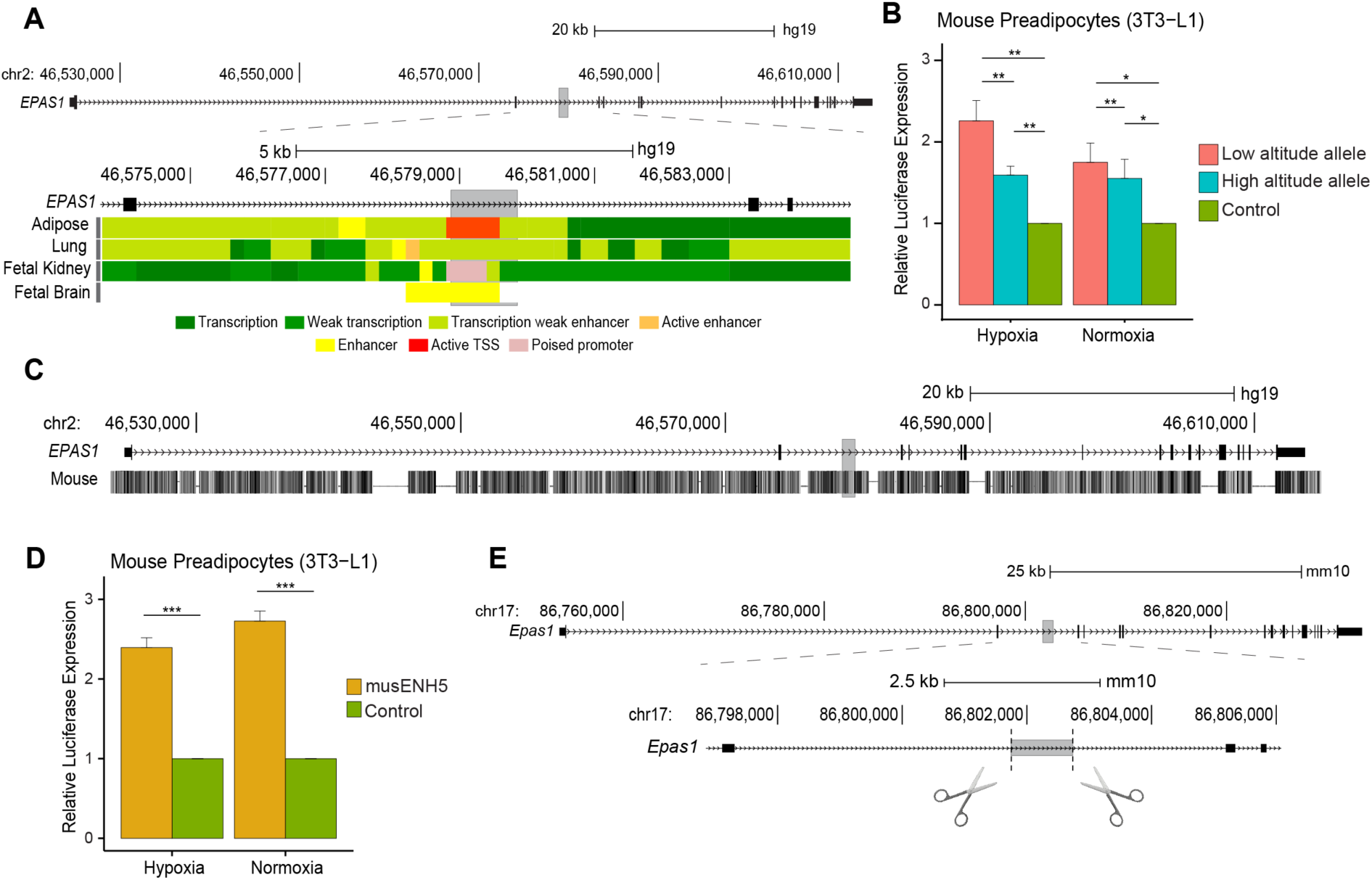
ENH5 is active in adipose tissue and is conserved in sequence and function in mouse adipocytes. (A) The genomic structure of the human *EPAS1* locus (chr2: 46524547-46613836, hg19). The locus is zoomed in at the ENH5 region (gray highlight: chr2: 46578867-46579856, hg19).Chromatin state tracks from the NIH Roadmap Epigenomics Mapping Consortium are shown for adipose, lung, fetal kidney, and fetal brain tissue. (B) Luciferase assay results of ENH5 low- and high-altitude alleles in 3T3-L1 cells in hypoxia (1% O_2_, 48 hours) and normoxia. The average ± SEM of 6 technical replicates (n=6) is shown. (C) The human *EPAS1* sequence is highly conserved in mice. The gray highlight indicates ENH5. (D) Luciferase assay results of mouse ENH5 in 3T3-L1 cells in hypoxia (1% O_2_, 48 hours) and normoxia. The average ± SEM of 6 technical replicates (n=6) is shown. (E) The genomic structure of the mouse *EPAS1* locus (chr17: 86753864-86833410, mm10). The locus is zoomed in at the ENH5 region (chr17: 86801737-86802738, mm10). The cut sites denote the target sites of the CRISPR gRNAs used to generate the homozygous ENH5 KO mouse line that has been previously established^9^. Data were analyzed using a one-tailed t-test. * p < 0.05, ** p < 0.01, and *** p< 0.001

### ENH5 does not impact adipose biology *in vivo* in the absence of cold or hypoxic stress

Having established that ENH5 has enhancer activity in mouse adipocytes under normoxic and hypoxic conditions, we first wanted to establish if ENH5 confers effects in adipocytes under normoxic conditions before elucidating any roles ENH5 might have under hypoxia or cold stress. We therefore assessed if ENH5 is conferring effects *in vivo* on adiposity and metabolic response in the absence of hypoxic or cold stress. To do so, we subjected male and female WT and ENH5 KO mice to a 12-week high fat diet (HFD) under standard mouse husbandry conditions, i.e. normoxia and room temperature (Figure 2A). We measured the weight of the mice weekly over the course of the HFD. We observed an increase in weight in both male and female mice during the HFD, but we did not observe any significant differences in weight between WT and ENH5 KO males or females (Figure 2A). Before and after the 12-week HFD, we measured abdominal fat percentage in male and female WT and ENH5 KO mice.Before the HFD, we did not observe a difference in abdominal fat percentage between WT or ENH5 KO male or female mice (Figure 2B). After the HFD, male and female mice gained abdominal fat, but we did not see a significant difference in abdominal fat gain between WT and ENH5 KO within either sex (Figure 2B). To assess if ENH5 has metabolic consequences *in vivo*, we performed intraperitoneal glucose tolerance tests (IPGTT) before and after the 12-week HFD. We did not observe any differences in glucose tolerance between WT and ENH5 KO mice in either sex before or after the HFD (Figure 2C). We also conducted histology on subcutaneous white fat post HFD to measure adipocyte number and size. We did not observe any significant differences in cell number or size between WT or ENH5 KO within each sex in the subcutaneous fat depots (Figure 2D). Taken together, these data indicate that, under normoxia and at room temperature, ENH5 does not have a detectable impact on adiposity or metabolic response in male or female mice *in vivo*.

**Figure 2.**
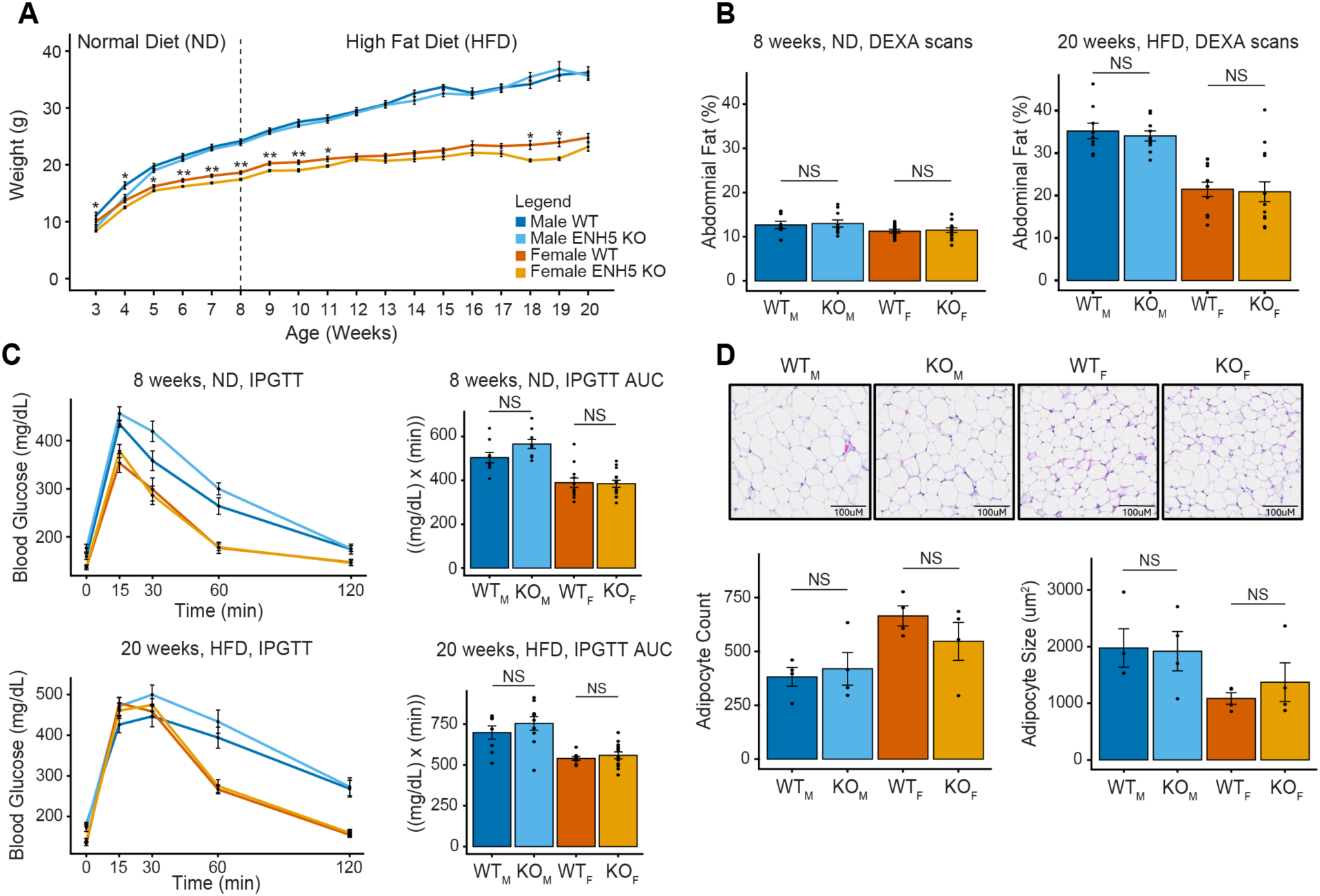
ENH5 does not impact adipose biology or metabolic response *in vivo*. (A) 8 week-old male and female WT and ENH5 KO mice were placed on a 12-week high-fat diet (HFD) (n = 9 male WT mice (WTM), n = 10 male ENH5 KO mice (KOM), n = 12 female WT mice (WTF) and n = 14 female ENH5 KO mice (KOF). Weight was measured weekly from weening (week 3) until the end of the HFD. (B) Abdominal fat percentage of mice before and after the HFD obtained from DEXA scans. (C) Intraperitoneal glucose tolerance tests (IPGTT) in WT and ENH5 KO mice before and after the HFD. Bar graphs represent area under curve (AUC). (D) Sections of subcutaneous fat from WT and ENH5 KO mice after the HFD. Bar graphs show cell count and cell size for 4 mice per genotype per sex. Data are presented as average ± SEM. Data were analyzed using two-tailed t-test. * p < 0.05 and ** p < 0.01.

### ENH5 regulates aerobic respiration and adipogenesis in adipocytes under thermogenic stimulation *in vitro*

Since we did not observe detectable effects on *in vivo* phenotypes in the absence of hypoxic and cold stress, we hypothesized that ENH5 may confer conditional effects dependent on environmental settings. To establish if ENH5 has measurable effects in adipocyte biology under hypoxic and cold stress we turned to an *in vitro* system. An important function of adipose tissue is to produce heat in cold temperatures by breaking down stored energy, which gets fed into the respiratory electron transport chain where it is uncoupled from making ATP and is released as heat^27–29^. Since Tibetans experience cold temperatures, we were motivated to assess if ENH5 KO impacts the thermogenic activity of adipocytes. To that end, we used an *in vitro* adipocyte differentiation model where we treated differentiated adipocytes with a β_3_-AR agonist drug used as a cold proxy. This drug, CL316243, is a β_3_-AR agonist that initiates thermogenic signaling via β_3_-AR activation. Specifically, we isolated preadipocytes from WT and ENH5 KO subcutaneous fat, which we then differentiated into mature adipocytes *in vitro*. We treated WT and ENH5 KO adipocytes in parallel with vehicle control or CL316243 for 4 hours (Figure 3A). After treatment, we collected the adipocytes and conducted RNA-seq and gene ontology (GO) analysis on genes that were differentially expressed in each of the genotypes in response to treatment. From previous studies, we anticipated that CL316243 treatment would induce changes in aerobic respiration and the electron transport chain^37–39^. In our data, we observed that both ENH5 KO and WT adipocytes downregulated pathways associated with aerobic respiration and the electron transport chain in response to treatment (Supplemental Figure 1A) [see supplemental text file]. Since both genotypes downregulate the same pathways, we wanted to know if the magnitude of the ENH5 KO response was different than the WT response, i.e. whether ENH5 KO responds to thermogenic stimulation with stronger, weaker, or the same level of downregulation as WT. To assess this, we used an interaction analysis for genotype x treatment (CL316243 treatment or vehicle treatment), then performed gene set enrichment analysis (GSEA) to identify pathways significantly enriched in differentially expressed genes in ENH5 KO vs WT adipocytes in response to treatment (Figure 3A). Interestingly, when we performed the interaction analysis for genotype x treatment, we found that ENH5 KO exhibited stronger differential expression of genes in the aerobic respiration and electron transport chain pathway compared to WT (Figure 3B). The genes that made up the aerobic respiration and respiratory electron transport gene set encode different subunits of the complexes that form the respiratory electron transport chain or encode proteins necessary for the TCA cycle, whose products feed into the respiratory electron transport chain (Figure 3C).

**Figure 3.**
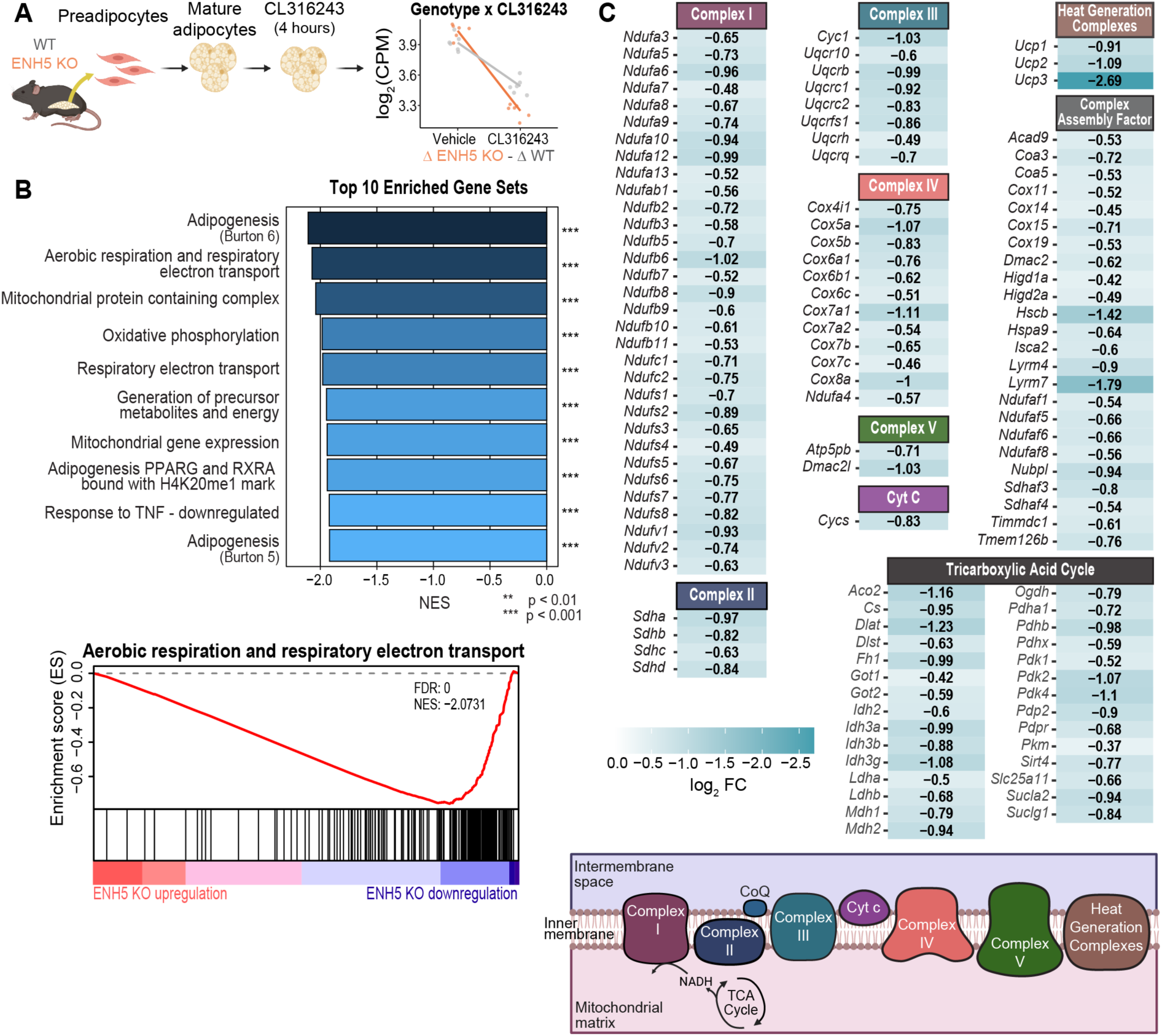
ENH5 KO more strongly downregulates expression of key genes involved in thermogenesis response upon CL316243 treatment compared to WT. (A) Experimental design showing preadipocytes being isolated from the subcutaneous fat pads of male WT (n = 6) and ENH5 KO mice (n = 6). Preadipocytes were then differentiated into mature adipocytes *in vitro* and stimulated with CL316243 for 4 hours. An interaction analysis for genotype by treatment was conducted to identify differentially expressed genes between WT and ENH5 KO response to treatment. (B) GSEA showing the top 10 enriched gene sets more strongly downregulated in ENH5 KO response than WT response to CL316243 treatment. The GSEA enrichment plot shows the enrichment score for the aerobic respiration and respiratory electron transport gene set. The red line indicates the running enrichment score (y-axis) while going down the ranked gene list (x-axis). The horizontal bars indicate the position of genes from the given gene set in the ranked gene list, where those at the top of the list are more upregulated in ENH5 KO response to CL316243 treatment compared to WT response while those at the bottom of the list are more downregulated in ENH5 KO response. Adipogenesis Burton 5 (MSigDB: M1564) and Adipogenesis Burton 6 (MSigDB: M1678) represent adipogenesis signaling pathways at early and late differentiation, respectively. (C) Heatmaps of the genes making up the aerobic respiration and respiratory electron transport gene set. Genes are clustered based on the electron complex chain component they are associated with.

Interestingly, we also observed that both WT and ENH5 KO adipocytes downregulated adipogenesis and associated PPAR signaling pathways (Supplemental Figure 1A). In our interaction analysis, we also saw adipogenesis as a top gene set where ENH5 KO exhibited stronger differential expression of genes in this pathway than WT in response to thermogenic stimulation (Supplemental Figure 1B). The genes that made up this pathway have roles in adipocyte differentiation, peroxisome function, mitochondrial function, and metabolism of fatty acids and lipids (Supplemental Figure 1C). Several key genes involved in these pathways were differentially downregulated in our dataset, such as *Adipoq*, *Cidec*, *Fasn*, *Plin4*, and *Slc2a4*, indicating these genes were more strongly downregulated in ENH5 KO adipocytes than in WT adipocytes in response to thermogenic stimulation. In aggregate, we found that ENH5 KO adipocytes responded to thermogenic stimulation with a stronger differential expression of key genes involved in the respiratory electron transport chain and adipogenesis compared to WT adipocytes.

### ENH5 regulates aerobic respiration and adipogenesis in adipocytes under hypoxia *in vitro*

Next, we set out to establish if ENH5 confers effects in adipocytes under hypoxic conditions. Though Tibetans experience cold temperatures seasonally, they are constantly in a hypoxic environment, thus motivating this study. Using the same *in vitro* adipocyte differentiation model used for cold stress, we differentiated WT and ENH5 KO preadipocytes into mature adipocytes and cultured them in hypoxia (1% O_2_) for 48 hours (Figure 4A). Again, we conducted RNA-seq and GO analysis on genes that were differentially expressed in each of the genotypes in response to hypoxia. Among our upregulated genes, we observed enrichment for response to hypoxia, as previously reported^40^, confirming our treatment was effective (Supplemental Figure 2A). We again expected to see changes in aerobic respiration and the electron transport chain pathways because hypoxic stress regulates these pathways in adipocytes to control oxygen use in metabolism^40,41^. As expected, we observed that both ENH5 KO and WT adipocytes downregulated pathways associated with aerobic respiration and the electron transport chain in response to hypoxia (Supplemental Figure 2B). Just as with the response to CL316243 treatment, we wanted to know if the magnitude of the ENH5 KO response was different than the WT response. To assess this, we used an interaction analysis for genotype x condition (hypoxia or normoxia), then performed GSEA (Figure 4A). Interestingly, when we performed the interaction analysis for genotype x condition, we found that ENH5 KO exhibited stronger differential expression of genes in the aerobic respiration and electron transport chain pathway compared to WT (Figure 4B). Similar to our results from the CL316243 treatment experiment, many of the genes in this gene set encode different subunits of the complexes that form the respiratory electron transport chain or encode proteins necessary for the TCA cycle, whose products feed into the respiratory electron transport chain (Figure 4C).

**Figure 4.**
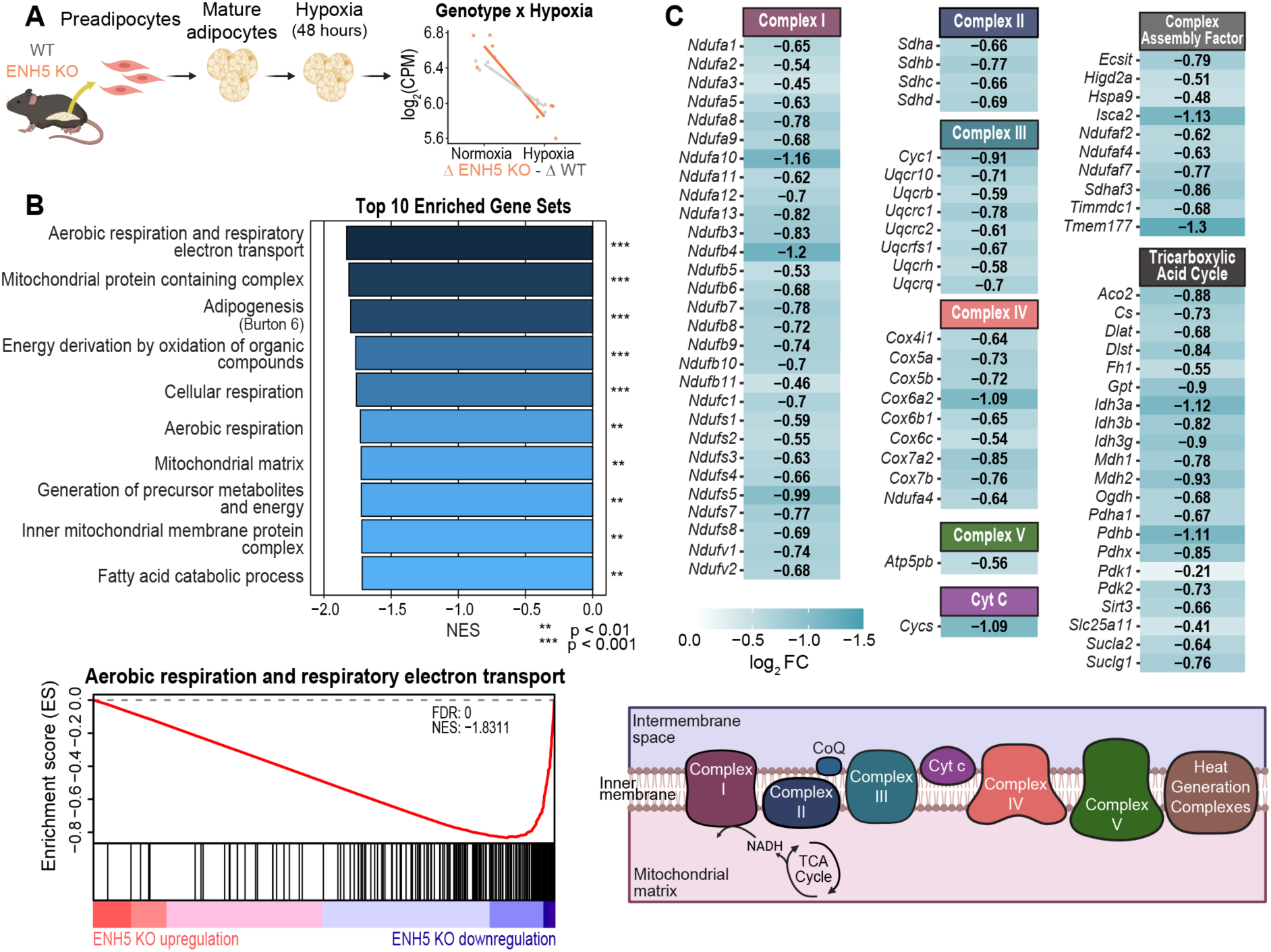
ENH5 KO more strongly downregulates expression of key genes involved in thermogenesis response under hypoxic conditions compared to WT. (A) Experimental design showing preadipocytes being isolated from the subcutaneous fat pads of male WT (n = 5) and ENH5 KO mice (n = 4). Preadipocytes were then differentiated *in vitro* to mature adipocytes, then cultured in hypoxia (1% O_2_) for 48 hours. An interaction analysis for genotype by treatment was conducted to identify differentially expressed genes between WT and ENH5 KO response to hypoxia. (B) GSEA showing the top 10 enriched gene sets more strongly downregulated in ENH5 KO response than WT response to hypoxia is shown. The GSEA enrichment plot shows the enrichment score for the aerobic respiration and respiratory electron transport gene set. The red line indicates the running enrichment score (y-axis) while going down the ranked gene list (x-axis). The horizontal bars indicate the position of genes from the given gene set in the ranked gene list, where those at the top of the list are more upregulated in ENH5 KO response to hypoxia compared to WT response while those at the bottom of the list are more downregulated in ENH5 KO response. Adipogenesis Burton 6 (MSigDB: M1678) represents adipogenesis signaling pathways at late differentiation. (C) Heatmaps of the genes making up the aerobic respiration and respiratory electron transport gene set. Genes are clustered based on the electron complex chain component they are associated with.

We also observed that both WT and ENH5 KO adipocytes downregulated the PPAR signaling pathway (Supplemental Figure 2B). In conjunction with these responses, we also saw adipogenesis as a top gene set in our interaction analysis where ENH5 KO exhibited stronger differential expression of genes in this pathway than WT in response to hypoxia (Supplemental Figure 2C). The genes in this pathway that were downregulated have roles in adipocyte differentiation, peroxisome function, mitochondrial function, and metabolism of fatty acids and lipids (Supplemental Figure 2D). Several key genes involved in these pathways were differentially downregulated in our dataset similar to what we observed in our thermogenic stimulation experiment, such as *Adipoq*, *Cidec*, *Fasn*, *Plin4*, and *Slc2a4*, indicating these genes were more strongly downregulated in ENH5 KO adipocytes than in WT adipocytes in response to hypoxia. Similar to what we observed with a thermogenic stimulus, we found that ENH5 KO adipocytes also responded to hypoxia with stronger differential expression of genes in two key pathways compared to WT adipocytes: the respiratory electron transport chain and adipogenesis. These data in total provide evidence for a role of ENH5 in regulating expression of the respiratory electron transport chain pathway and the adipogenesis pathway in response to thermogenic stimulation and hypoxic conditions independently.

### Combined thermogenic stimulation and hypoxia stress does not result in an additive response in ENH5 KO adipocytes

Having established that ENH5 KO adipocytes exhibit stronger differential expression of genes in aerobic respiration and adipogenesis pathways compared to WT adipocytes in response to thermogenic stimulation and hypoxia independently, we wanted to assess how ENH5 KO adipocytes responded to these stresses together. Tibetans are always experiencing hypoxia, but only experience cold stress seasonally. When experiencing cold stress in an already hypoxic environment, perhaps ENH5 KO responds to this joint condition with an additive effect, meaning that adding cold stress would result in ENH5 KO exhibiting an even stronger differential expression of genes in aerobic respiration and adipogenesis pathways. To test this, we assessed how ENH5 KO adipocytes respond to the addition of thermogenic stimulation in a hypoxic background. Using the same *in vitro* adipocyte differentiation model, we first cultured mature adipocytes in hypoxia (1% O^2^) for 48 hours. We subsequently treated the adipocytes with CL316243 for an additional 4 hours in hypoxia and conducted RNA-seq to identify differential response between hypoxia and hypoxia + CL316243 conditions (Supplemental Figure 3A). If ENH5 KO were to exhibit an additive effect on aerobic respiration and adipogenesis pathways, we would first expect to see these pathways enriched in downregulated DEGs in the individual WT and ENH5 KO response, meaning that these pathways are being downregulated when thermogenic stimulation is added to a hypoxic background. Secondly, we would expect to see enrichment of these pathways in our interaction analysis, meaning that the response of ENH5 KO relative to WT differs in the combined versus the individual treatments. From our GO analysis, we observed that WT and ENH5 KO adipocytes downregulate adipogenesis, but not aerobic respiration, in response to CL316243 treatment under hypoxia (Supplemental Figure 3B). To determine if the magnitude of the ENH5 KO response was different than the WT response, we used an interaction analysis for genotype x treatment (hypoxia + CL316243 treatment or hypoxia + vehicle treatment), then performed GSEA (Supplemental Figure 3A). We did not find any gene sets to be significantly enriched for differential response to thermogenic stimulation in a hypoxic background between ENH5 KO and WT adipocytes (Supplemental Figure 3C). While we observed that both WT and ENH5 KO downregulated pathways in response to CL316243 treatment in a hypoxic background, these pathways were not being differentially downregulated between the genotypes (Supplemental Figure 3D). Therefore, the combination of thermogenic stimulation in a hypoxic environment does not confer a detectable additive response in ENH5 KO adipocytes compared to WT adipocytes. In conclusion, though we showed that ENH5 KO adipocytes have stronger differential expression of aerobic respiration and adipogenesis genes compared to WT adipocytes in response to thermogenic stimulation and hypoxia independently, we did not observe an additive effect of ENH5 KO response when treated with both conditions simultaneously.

## DISCUSSION

In this work, we characterize the functional role of ENH5 at the *EPAS1* locus in subcutaneous white adipocytes. We show the Tibetan ENH5 haplotype is active in adipose tissue, displaying allele-specific effects compared to the lowlander haplotype. Dissecting the role of ENH5 using an ENH5 KO mouse model, our data suggests that ENH5 does not impact fat composition or metabolism *in vivo* under standard animal husbandry conditions, but it does have conditional effects under thermogenic stimulation and hypoxia independently *in vitro*. Under either of these conditions, ENH5 KO adipocytes display stronger differential expression of genes in aerobic respiration and adipogenesis pathways compared to WT adipocytes. Yet, under these conditions jointly, we do not observe an additive response in ENH5 KO adipocytes compared to WT. Our results expand on the pleiotropic role of ENH5 in two ways: 1) by showing it has effects in adipocytes and 2) by showing ENH5 responds to thermogenic stimulation in addition to responding to hypoxia.

We show that ENH5 is an active enhancer in adipocytes with allele-specific effects, where the high-altitude allele has reduced enhancer activity compared to the low-altitude allele (Figure 1B). These allele-specific effects of ENH5 in adipocytes are consistent with previous work establishing that the Tibetan ENH5 allele has reduced enhancer activity in multiple hypoxia-responsive cell types^42^. Here, we have established that ENH5 is active in yet another tissue type, further characterizing the pleiotropic nature of this enhancer and contributing to the growing evidence of enhancers having pleiotropic roles across different tissue types^43–45^.

Using an ENH5 KO mouse model, we show that ENH5 KO in adipocytes not only has a differential response to hypoxia, but thermogenic stimulation as well. Previous work has established that ENH5 KO confers a differential response compared to WT under hypoxia, where ENH5 KO mice exposed to hypoxia were shown to have differential gene expression in several tissue types, including the lung, right atrium, and left ventricle^42^. Our work shows another environmental condition where ENH5 KO confers differential response: thermogenic stimulation. We show that both WT and ENH5 KO adipocytes respond to hypoxia and thermogenic stimulation by downregulating aerobic respiration and adipogenesis pathways (Figures 3 and 4). While we expected both genotypes to downregulate these pathways in response to hypoxia based on previous work^40,41^, we were surprised to see this response in both genotypes in response to CL316243 treatment. While this downregulation in response to thermogenic stimulation may be due in part to a local hypoxic environment of our *in vitro* model (see supplemental text), we observe that ENH5 KO adipocytes have stronger differential expression of genes in these pathways compared to WT adipocytes, demonstrating a difference in response due to genotype. This local hypoxic environment is a limitation of our model system and future *in vivo* work is needed to disentangle the response of ENH5 KO to thermogenic stimulation versus hypoxia.

Under thermogenic stimulation and hypoxia independently, we observe a difference in transcriptional responses between ENH5 KO and WT adipocytes. Our model was only able to assess transcriptional responses, since the β_3_-AR agonist drug activates thermogenic signaling and is not capturing the full physiological response to cold stress response, which also involves shivering and vasoconstriction^46^. For aerobic respiration, we observe key OXPHOS and TCA cycle genes being more strongly differentially expressed in ENH5 KO adipocytes compared to WT adipocytes. The stronger differential expression of these genes means that the aerobic respiration pathway is being suppressed more in ENH5 KO adipocytes, suggesting that less energy is being fed into the electron transport chain to be released as heat. Similarly, we observe that key genes involved in adipogenesis are more strongly differentially expressed in ENH5 KO response to these conditions compared to WT. In response to thermogenic stimulation, new adipocytes can be generated to respond to the thermogenic demand^47–49^. The stronger downregulation of the adipogenesis pathway we see suggests that fewer adipocytes are being generated to respond to thermogenic stimulation. While future studies need to be conducted especially *in vivo* to understand the full physiological response to cold stress, the stronger differential expression of genes in these pathways in ENH5 KO adipocytes suggests that ENH5 KO is more strongly suppressing thermogenesis. This response may be conferring lower thermogenic activity in response to these environmental conditions in Tibetans, which may act to conserve energy instead of expending it as heat.

Our findings that ENH5 KO regulates thermogenic activity in adipocytes is supported in the literature. HIF-2α, the protein product of *EPAS1*, has been implicated as a negative regulator of thermogenic response to protect against exhaustive energy expenditure^33^. While we have clearly demonstrated that ENH5 is involved in the regulation of thermogenic activity, how ENH5 regulates *EPAS1* expression and subsequent protein levels of HIF-2α to regulate thermogenic activity is unclear. *EPAS1* expression and HIF-2α protein levels are dynamic in response to hypoxia^50^. Additionally, this dynamic *EPAS1* expression is still observed with deletion of ENH5, possibly due to several enhancers shown to have regulatory activity at this locus^42^. While our study lacks a time course to interrogate this dynamic expression, future studies incorporating multiple treatment time points to capture the dynamic expression of *EPAS1* and protein levels of HIF-2α in response to CL316243 treatment and hypoxia will need to be conducted to understand the mechanism of how ENH5 regulates thermogenic activity.

To contextualize our results with regards to the Tibetan ENH5 haplotype, our data postulate that the Tibetan ENH5 haplotype is exhibiting stronger downregulation of thermogenic activity to hypoxia and thermogenic stimulation, independently. Since the ENH5 KO mouse phenocopies the Tibetan ENH5 haplotype which has reduced enhancer activity, our data suggests that Tibetans could have lower thermogenic activity in response to these conditions, independently. Though Tibetans experience cold stress and hypoxia jointly, and our data do not show ENH5 KO to have an additive response to these conditions, our results suggest that ENH5 KO may have a role in regulating thermogenesis. One limitation in our work is that we might be missing a differential response between genotypes under the joint conditions since we did not assess chronic stress conditions. Future work with longer treatment time points will be more representative of the environmental stressors that Tibetans experience and will help elucidate the role of ENH5 KO under chronic stress. Additionally, the extrapolation of our results to a Tibetan phenotype are speculative and need to be confirmed with future studies. *In vivo* work using ENH5 KO mice under chronic stress will establish what organismal phenotypes are being impacted and can be assessed to understand how these phenotypes may be impacting fitness. Additionally, a human iPSC model established from Tibetans provides a great human model to study the effects of the Tibetan ENH5 haplotype^10^. Though our data was conducted *in vitro*, our work provides strong evidence that ENH5 KO has a role in thermogenic regulation in adipocytes, and warrants future studies to assess the possible role the Tibetan ENH5 haplotype might have on thermogenic activity and balancing heat generation and energy expenditure under hypoxic and cold stress.

In support of motivating future work to assess the role of the Tibetan ENH5 haplotype in thermogenic response, thermogenic response has been shown to be an adaptive phenotype for mammals living at high altitude and experiencing cold temperatures. For example, deer mice living at higher altitudes were found to have more upregulation of OXPHOS genes compared to deer mice at lower altitudes^51^. Additionally, plateau pika living in the Tibetan Plateau have been shown to have increased thermogenic properties, including increased expression of thermogenic genes and increased transitioning of white adipose tissue to brown adipose tissue^52^. Furthermore, another study in plateau pika found thermogenic genes under positive selection in high-altitude pika compared to low-altitude pika^53^. Previous studies have shown that all these species, like humans, rely on genetic variants in the *EPAS1* locus for their adaptation in the hypoxic and cold Tibetan Plateau, lending support to our findings that thermogenesis is a pathway that is being regulated by HIF-2α in response to cold stress and high altitude. While these studies in small mammals show that upregulation of thermogenesis is an adaptive phenotype, our work suggests that the Tibetan ENH5 haplotype confers downregulation of thermogenesis in response to these environmental stressors. A possible explanation for this difference is that small mammals rely more on thermogenesis to maintain body temperature, while humans have other methods, such as clothing, to regulate body temperature. Small mammals experience excessive heat loss compared to humans based on their surface-to-volume ratio, and thus rely heavily on thermogenesis from brown adipocytes, which is a specific type of adipocyte whose predominant function is to generate heat^54^. Humans, on the other hand, rely on brown adipocytes at birth to regulate body temperature, but quickly lose this type of adipocytes thereafter^30,31^. It is possible that the downregulation of thermogenesis genes we observed in the ENH5 KO reflect the ability of Tibetans to buffer themselves from cold temperatures. Tibetans can rely more on behavioral thermoregulation (e.g. clothing, shelter, fire) than physiological thermogenesis, making energy conservation more beneficial than heat production. For example, Tibetans wear traditional robes that are typically thick and can be worn in different ways to adjust their thermal insulation^55,56^. Since Tibetans can physically adapt to cold temperatures through their clothing and behavior, it might be beneficial for Tibetans to downregulate thermogenesis to decrease energy expenditure that would be used to produce unnecessary heat to maintain body temperature. Our results suggest that the Tibetan ENH5 haplotype may favor energy storage rather than energy expenditure under cold and hypoxic conditions.

In summary, we demonstrate a further pleiotropic role of ENH5, where this enhancer is active in adipocytes and regulates thermogenic activity in response to thermogenic stimulation and hypoxia, independently. We establish that adipocytes may be an important cell type to study when dissecting Tibetan adaptation to high altitude. Additionally, we show that cold stress may be an important environmental condition that is affecting Tibetan adaptation. While further work needs to be conducted to fully establish the role of the Tibetan ENH5 haplotype in thermogenic regulation, our work suggests that ENH5 regulation of thermogenic activity may be under selection to balance energy expenditure and heat generation in cold stress at high altitudes. Our findings provide a possible physiological explanation for why the *EPAS1* locus is under such strong selection in humans and mammals in the Tibetan Plateau.

---

**Supplemental Figure 1.**
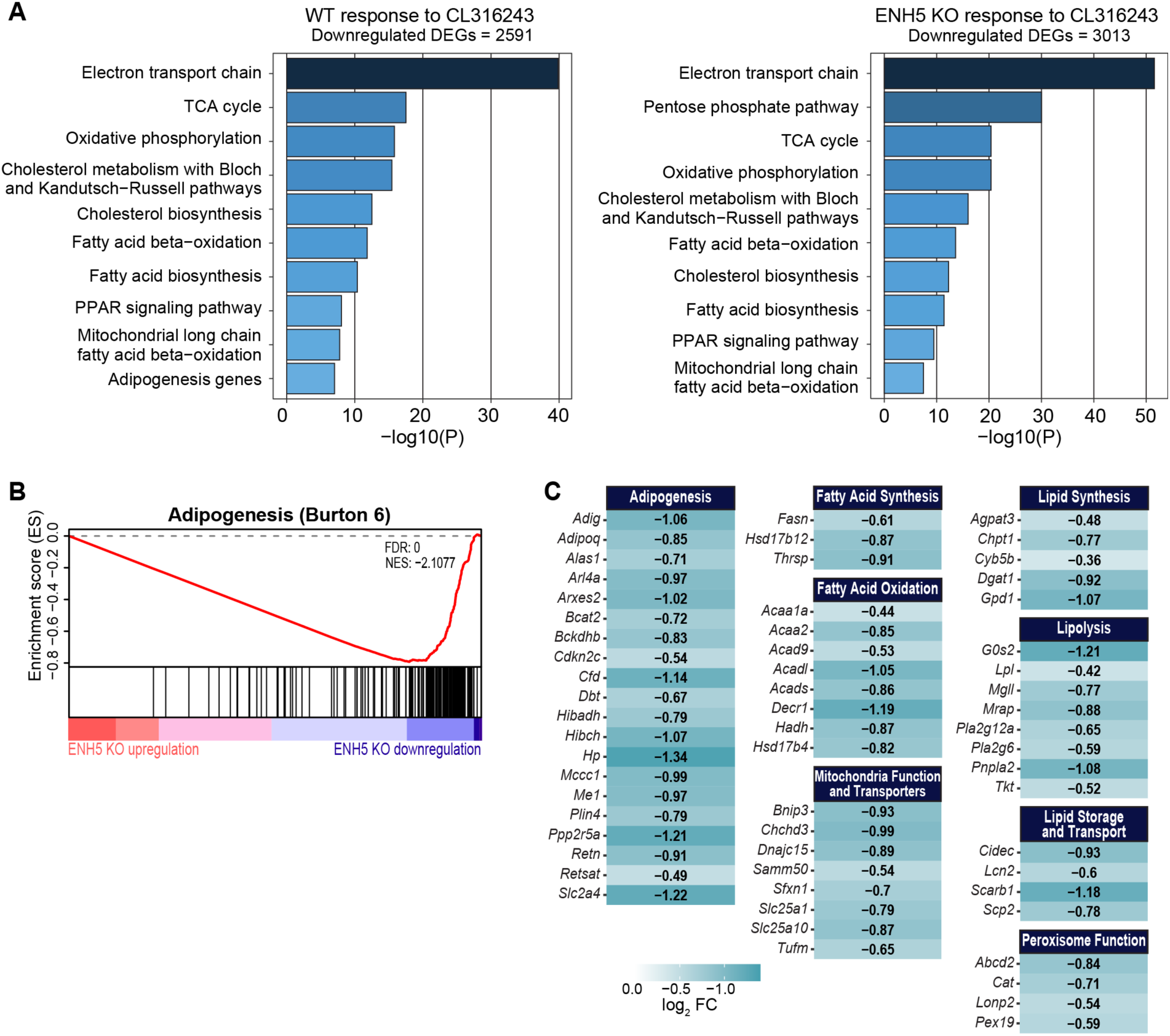
WT and ENH5 KO response to CL316243 treatment. (A) Top 10 enriched pathways from WikiPathways for WT and ENH5 KO genes downregulated in response to CL316243. n= 6 for male WT mice and n = 6 for male ENH5 KO mice. (B) GSEA enrichment plot showing the enrichment score for the adipogenesis gene set. The red line indicates the running enrichment score (y-axis) while going down the ranked gene list (x-axis). The horizontal bars indicate the position of genes from the given gene set in the ranked gene list, where those at the top of the list are more upregulated in ENH5 KO response to CL316243 treatment compared to WT response while those at the bottom of the list are more downregulated in ENH5 KO response. Adipogenesis Burton 6 (MSigDB: M1678) represents adipogenesis signaling pathways at late differentiation. (C) Heatmaps of the genes making up the adipogenesis gene set. Genes are clustered based on different roles of adipogenesis they are associated with.

**Supplemental Figure 2.**
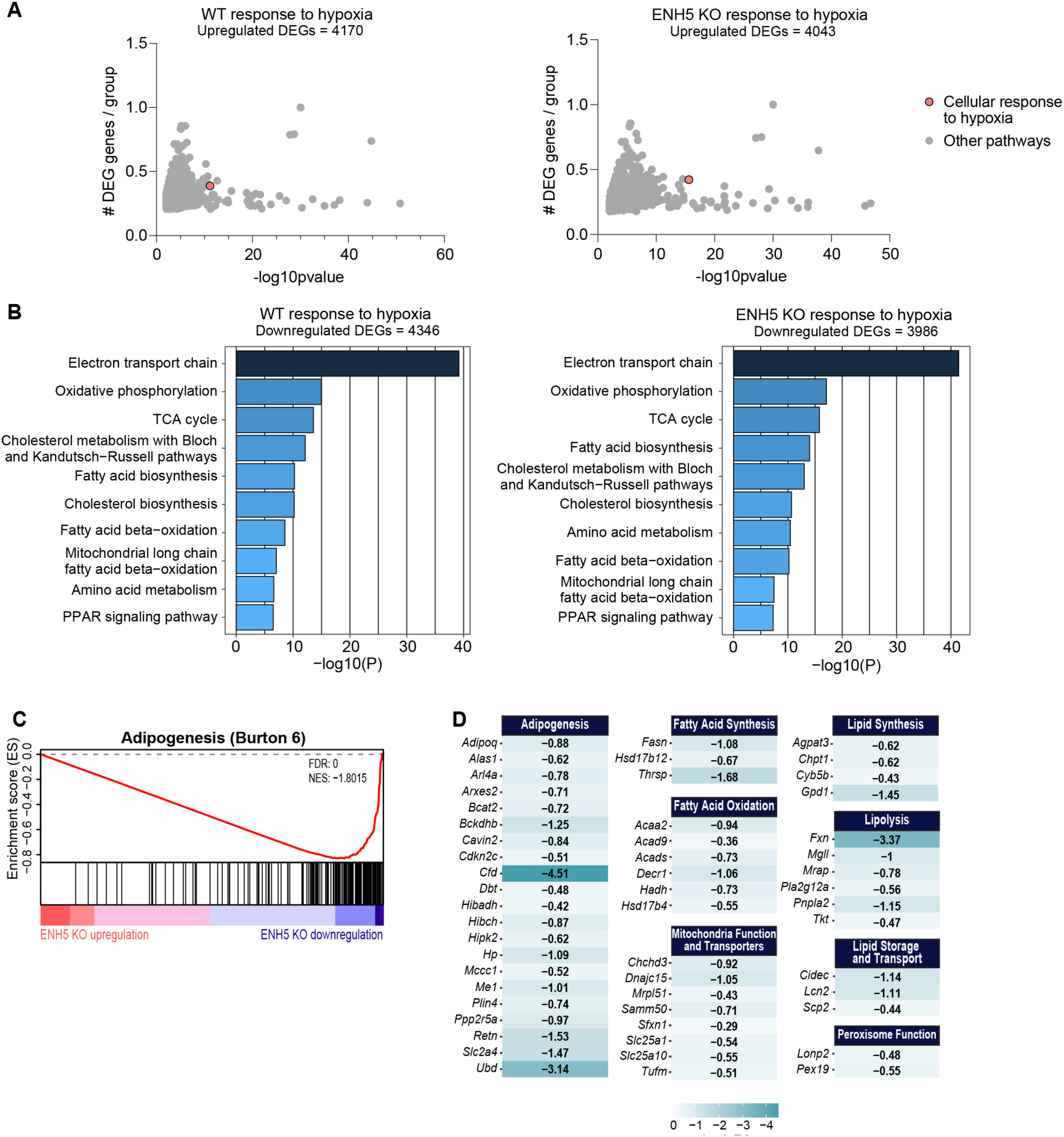
WT and ENH5 KO response to hypoxia. (A) Biological processes enrichment using gene ontology analysis of WT and ENH5 KO genes upregulated in response to hypoxia. n = 5 for male WT mice and n = 4 for male ENH5 KO mice. (B) Top 10 enriched pathways from WikiPathways for WT and ENH5 KO genes downregulated in response to hypoxia. (C) GSEA enrichment plot showing the enrichment score for the adipogenesis gene set. The red line indicates the running enrichment score (y-axis) while going down the ranked gene list (x-axis). The horizontal bars indicate the position of genes from the given gene set in the ranked gene list, where those at the top of the list are more upregulated in ENH5 KO response to hypoxia compared to WT response while those at the bottom of the list are more downregulated in ENH5 KO response. Adipogenesis Burton 6 (MSigDB: M1678) represents adipogenesis signaling pathways at late differentiation. (D) Heatmaps of the genes making up the adipogenesis gene set. Genes are clustered based on different roles of adipogenesis they are associated with.

**Supplemental Figure 3.**
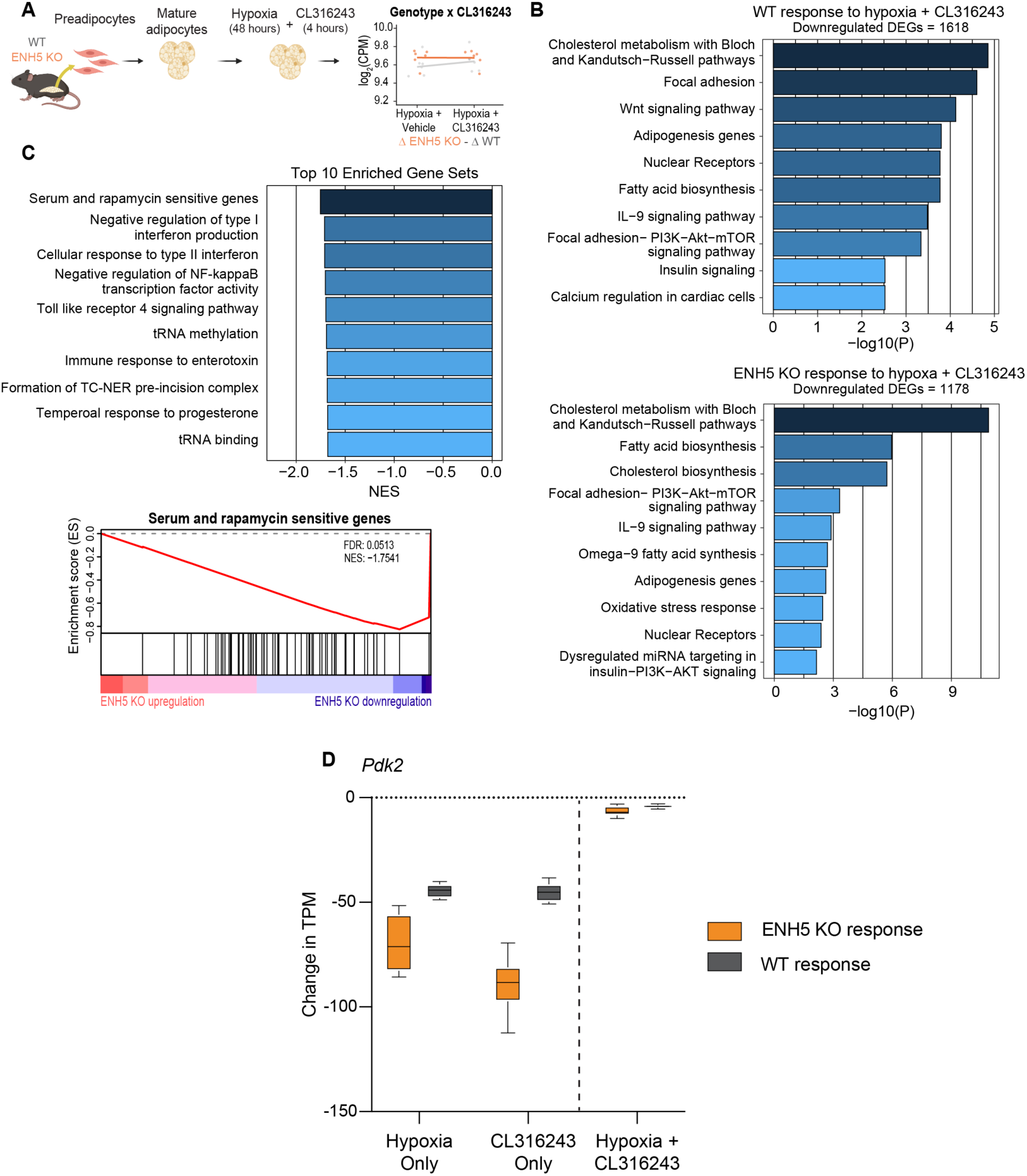
Joint hypoxia and thermogenic stimulation does not have an additive effect on transcriptional regulation in ENH5 KO adipocytes. (A) Experimental design showing preadipocytes being isolated from the subcutaneous fat pads of male WT (n = 6) and ENH5 KO mice (n = 6). Preadipocytes were differentiated *in vitro* to mature adipocytes, then cultured in hypoxia (1% O2) for 48 hours followed by a 4 hour CL316243 treatment while in hypoxia. An interaction analysis for genotype by treatment was conducted to identify differentially expressed genes between WT and ENH5 KO response to thermogenesis in hypoxia. (B) Top 10 enriched pathways from WikiPathways for WT and ENH5 KO genes downregulated in response to hypoxia + CL316243. (C) GSEA showing the top 10 enriched gene sets more strongly downregulated in ENH5 KO response than WT response to the joint treatment is shown. The GSEA enrichment plot shows the enrichment score for the serum and rapamycin sensitive genes gene set. The red line indicates the running enrichment score (y-axis) while going down the ranked gene list (x-axis). The horizontal bars indicate the position of genes from the given gene set in the ranked gene list, where those at the top of the list are upregulated in ENH5 KO response to thermogenesis in hypoxia compared to WT response while those at the bottom of the list are downregulated in ENH5 KO response. (D) Box plot of the change in expression of *Pdk2* in ENH5 KO and WT adipocytes in response to all three experimental conditions. Dotted horizontal line separates hypoxia + CL16243 treatment to indicate that this experiment was conducted in a hypoxic background.

## ACKNOWLEDGEMENTS

We would like to acknowledge support from the staff at the following University of Chicago facilities: the Animal Resources Center Core Facility (RRID:SCR_021806), the Human Tissue Resource Center (RRID:SCR_019199), the Functional Genomics Core Facility (RRID:SCR_019196) and the Research Computing Cluster. This work was funded by the National Institutes of Health grants R01HL119577 (MAN and AD), 5T32GM139782-02 (AT), T32HL007605 (IMS), and T32HL007381 (ZTW) and the University of Chicago Diabetes Research Center grant P30 DK020595 (SP). Diagrams in Figures 3, 4, and Supplemental Figure 3 were generated with Biorender.

## METHODS

### Luciferase assays

Luciferase reporter assays for high- and low-altitude ENH5 alleles and mouse ENH5 allele were performed in 3T3-L1 cells as previously described^42^. 3T3-L1 cells were seeded in 96-well plates at 5,000 cells per well. 24 hours after plating when cells were at approximately 50% confluency, 3T3-L1 cells were co-transfected with 100ng of each construct and 10ng of pGL4.73[hRluc/SV40] *Renilla* luciferase reporter vector (Promega, E6911) to normalize transfection efficiency. 3T3-L1 cells were also transfected with a negative control construct (scramble) using a DNA sequence (chr9:6,161,550–6,162,024) devoid of any active chromatin epigenetic marks. Co-transfection of each construct occurred in triplicate using Lipofectamine LTX with Plus Reagent (Thermo Fisher Scientific, 153308030). After transfection, cells were cultured in normoxia or hypoxia (1% O^2^) for 48 hours. Luciferase activity was determined by measuring firefly and *Renilla* luciferase activity using the Dual-Luciferase Reporter Assay System (Promega, E1960) Briefly, cells were washed with PBS, lysed with 1X passive lysis buffer on a rocking platform for 15 minutes at room temperature, and stored at −20 °C until luciferase activity was measured. For all three measurements of each construct and the control, firefly luciferase activity was normalized to *Renilla* luciferase activity and the average luciferase/*Renilla* ratio was calculated (luciferase/*Renilla* will now be referred to as luciferase activity). The Dual-Luciferase assay was conducted four separate times using different DNA Lipofectamine transfection preparations where the cells were transfected, collected, and luciferase activity was measured on different days.

### Mice

All mouse procedures were approved by the Institutional Animal Care and Use Committee (IACUC) of the University of Chicago. ENH5 KO mice (strain C57BL/6J) were previously established^42^. Mice were housed on a 12 hour light/dark cycle with ad libitum access to food and water according to their assigned normal diet or high fat diet (HFD). For the high fat diet studies, 8-week-old male and female mice were placed on a 55% HFD (Envigo Teklad, TD.93075) for 12 weeks. Body weight was measured weekly from 3 to 20 weeks of age. Before and after HFD, abdominal fat mass was quantified by performing Dual-Energy X-ray Absorptiometry (DEXA) scans using a Lunar PIXImus II (GE Medical Systems). Abdominal fat mass was quantified using the Lunar PIXImus II analysis software by defining a region of interest from the manubrium (top of rib cage) to the bottom of the hip joint of the mouse.

### Metabolic phenotyping experiments

*In vivo* intraperitoneal glucose tolerance tests (IPGTT) were conducted in mice before and after the high fat diet. Mice were fasted for 4 hours, then injected intraperitoneally with dextrose (2g per kg body weight). Blood glucose levels were measured 0, 15, 30, 60, and 120 minutes post dextrose injection using a (Accu-Chek Aviva). The area under the curve (AUC) describing blood glucose levels post dextrose injection were calculated with the trapezoidal rule: AUC = 0.25 x (0 min fasting measurement) + 0.5 x (30 minute measurement) + 0.75 x (60 minute measurement) + 0.5 x (120 minute measurement). Two-sided t-tests between WT and ENH5 KO mice within each sex were conducted on AUC values to determine statistically significant difference in blood glucose levels with a significant p-value threshold of p < 0.05. Histology analysis was conducted on fat pads to quantify cell number and cell size after a 12-week HFD. After the 12-week HFD, subcutaneous fat pads were harvested from male and female WT and ENH5 KO mice. Fats pads were fixed in 4% formalin solution (Millipore Sigma, HT501128), embedded in paraffin, cut into 5μm sections, and stained with hematoxylin and eosin (H&E). Whole histology slides were scanned at the University of Chicago Integrated Light Microscopy Core using the Olympus VS200 Slideview Research Slide Scanner. Adipocyte size and number were quantified using Montpellier Resources Imagerie Adipocyte Tool^57^ with ImageJ software^58^. For 4 mice per genotype per sex, two H&E stained histology slides were analyzed per mouse where 3 randomly selected fields at 10X per slide were used for adipocyte quantification to get 6 measurements per mouse.

### *In vitro* differentiation

Primary stromal vascular cells from subcutaneous fat pads of 9 to 11 week old WT and ENH5 KO mice was fractioned, cultured, and differentiated as described^59^. Briefly, preadipocytes were isolated from the stromal vascular section from subcutaneous fat pads and were cultured in DMEM supplemented with 20% FBS, 40μg/mL penicillin-streptomycin, 40μg/mL gentamicin, and 500ng/mL amphotericin B. Once preadipocytes were at 90% confluency, cells were cultured in induction medium for 2 days, which consisted of DMEM supplemented with 10% FBS, 40μg/mL penicillin-streptomycin, 40μg/mL gentamicin, 850nM insulin, 1nM triiodothyronine (T3), 1μM rosiglitazone, 1μM dexamethasone, 500μM 3-isobutyl-1-methylxanthine (IBMX), and 125μM indomethacine. Cells were then cultured in differentiation media for 7 days, consisting of DMEM supplemented with 10% FBS, 40μg/mL penicillin-streptomycin, 40μg/mL gentamicin, 850nM insulin, 1nM triiodothyronine (T3), and 1μM rosiglitazone.Differentiation media was changed every 2 days over the course of the 7 day differentiation. At the end of the 7 day differentiation, the mature adipocytes were treated based on the specified experiment, then collected for RNA-sequencing. For hypoxia conditions, mature adipocytes were moved to a humidified incubator contained within a hypoxia chamber maintained at 37°C, 5% CO^2^, and 1% O^2^ for 48 hours. For thermogenic conditions, mature adipocytes were cultured for 4 hours in the presence or absence of 5μM CL316243 (Sigma, C5976). For the combined hypoxia and thermogenic conditions, mature adipocytes were cultured in the hypoxia chamber for 48 hours, then an additional 4 hours in hypoxia in the presence or absence of 5μM CL316243.

### RNA-sequencing

Total RNA was isolated from snap-frozen cell pellets using Qiagen RNeasy Mini kit (Qiagen, 74104). RNA quality was assessed using an Agilent Bioanalzyer. The RNA integrity number (RIN) for all samples was > 9. 1ug of RNA was used to prepare RNA-sequencing libraries using the NEBNext Ultra II Directional RNA Library Prep Kit for Illumina (NEB, E7765). Sequencing was performed at the University of Chicago Genomics Core Facility using an Illumina NovaSeq using 50bp paired-end sequencing.

### Differential gene expression

RNA-sequencing reads were aligned using STAR version 2.6.1b^60^ to the GRCm39 release 108. Any lowly expressed genes with < 1 cpm across most samples were filtered out of the analysis. edgeR version 4.4.2^61^ was used to normalize the read counts using mean of M-values (TMM) and logCPM.

Differential gene expression analysis for individual genotype response to experimental condition was conducted using edgeR version 4.4.2^61^ using the quasi-likelihood generalized linear model functions glmQLFit() and glmQLFTest() and linear model (∼0 + mouse + condition) for WT and ENH5 KO separately. Our model included mouse pairing in each experiment since mature adipocytes from each mouse were treated with the appropriate control and experimental condition. FDR was used for multiple testing correction and an FDR cutoff of 0.05 was used to determine significantly differentially expressed genes. Gene Ontology^62^ terms and gene associations were downloaded from https://geneontology.org on February 23, 2026. Enrichment was assessed by counting the number of terms annotated to a given gene set and to the universe of annotated genes. A chi-square test was performed to assess statistical significance (adjusted p < 0.01, fold-enrichment > 2 and > 5 genes).

Differential gene expression analysis using an interaction effect between genotype and experimental condition was conducted using a linear mixed-effects model using lme4 version 1.1.37^63^. The mixed-effects model accounted for mouse pairing in each experiment since mature adipocytes from each mouse were treated with the appropriate control and experimental condition (∼(1|mouse) + genotype*condition). P-values were adjusted for multiple test correction using FDR. Gene set enrichment analysis (GSEA)^64,65^ was used to determine genes with significant differential expression between WT and ENH5 KO response to the given experimental condition. GSEA analyses were performed with GSEAPreranked version 4.1.0 We ranked genes by multiplying the -1 X natural log adjusted p-value by the log2 fold-change of gene expression comparisons. We used -scoring_scheme weighted, -nperm 1000, -collpase No_Collapse, -norm meandiv, -set_min 15 and -set_max 500.

### Statistical analysis

Statistical analyses were performed using RStudio. For luciferase assays, the average luciferase activity for each construct was compared to the average luciferase activity for the control using a one-tailed paired t-test. For DEXA scan data, IPGTT, and histology analyses, a two-sided Student’s t-test was performed to determine significant differences between WT and ENH5 KO mice within each sex. A p-value threshold of 0.05 was used to determine significance.

### Data availability

All RNA-seq data generated in this work are available at Array Express (https://www.ebi.ac.uk/biostudies/) under accession number E-MTAB-17112.

