## Supplemental Text for "A pleiotropic *EPAS1* enhancer mediating Tibetan adaptation to hypoxia is active in adipocytes"

**Supplemental text for CL316243 treatment experiment**

We expected that WT and ENH5 KO adipocytes would individually respond to CL316243 treatment by upregulating aerobic respiration and the electron transport chain pathways since $\beta_3$ adrenergic receptor ($\beta_3$-AR) activation upregulates these pathways to release stored energy as heat[1–3]. However, in our experimental system, we observed that WT and ENH5 KO adipocytes individually responded to this treatment by downregulating aerobic respiration and the electron transport chain. We looked at the expression levels of *Adrb3*, the gene that encodes $\beta_3$-AR, and found that *Adrb3* was downregulated in response to CL316243 treatment, suggesting that we measured WT and ENH5 KO response to cold stress during a negative feedback loop that may result in lower thermogenic activity despite using a dosage and time point for CL316243 treatment that is effective at stimulating thermogenesis activity in other systems[4–6] (Supplemental Figure 1A). $\beta_3$-AR can be desensitized during thermogenic stimulation[7], and has been shown to be desensitized in mice as early as 2 hours after treatment[8]. Additionally, CL316243 treatment and hypoxia can indirectly cause hypoxia and a metabolic response, respectively: CL316243 treatment can cause local hypoxia since thermogenesis is an oxygen-dependent process[9,10], while hypoxia can downregulate aerobic respiration to regulate oxygen consumption in metabolic response[11,12]. To determine whether this was true also in our experimental system, we looked at key gene markers of hypoxia in adipocytes, and we found that several of these markers were upregulated in response to CL316243 treatment (Supplemental Figure 1B). Additionally, we found that cellular response to hypoxia is an enriched pathway among upregulated genes in both WT and ENH5 KO adipocytes in response to CL316243 (Supplemental Figure 1C). This suggested that this treatment caused a local hypoxic response in these adipocytes, thus causing the negative feedback loop exhibited by downregulation of $\beta_3$-AR. Additionally, for many of these hypoxic marker genes, we observed that the adipocytes used for the CL316243 treatment started at a higher expression before treatment, suggesting that the adipocytes might have experienced a hypoxic environment. As adipocytes expand in size and number, this can result in a local hypoxic environment[13]. Our *in vitro* model results in high adipocyte differentiation efficiency, which is a plausible explanation as to why the adipocytes used in this experiment exhibited hypoxia signatures before CL316243 treatment (Supplemental Figure 1D). While our model does not show the results we would expect to observe at other time points after treatment, and is thus a limitation of our model, understanding cellular response to the negative feedback loop that regulates thermogenesis is still an important component of cold stress response and is informative to understand how adipocytes respond to cold stress. In response to CL316243 treatment, we observed that WT and ENH5 KO adipocytes downregulated thermogenic pathways, but ENH5 KO exhibited stronger downregulation of these pathways compared to WT (Main Text Figure 3B).

**Supplemental Figure 1.** CL316243-treated adipocytes downregulate genes involved in thermogenic activity likely due to local hypoxic environment. (A) Expression changes of *Adrb3* in adipocytes treated independently with hypoxia or CL316243. (B) Expression changes of key hypoxia marker genes in adipocytes treated independently with hypoxia or CL316243. (C) Biological processes enrichment using gene ontology analysis of WT and ENH5 KO genes upregulated in response to hypoxia. (D) Images of differentiated WT and ENH5 KO adipocytes before CL316243 treatment. Scale bars: 400uM.

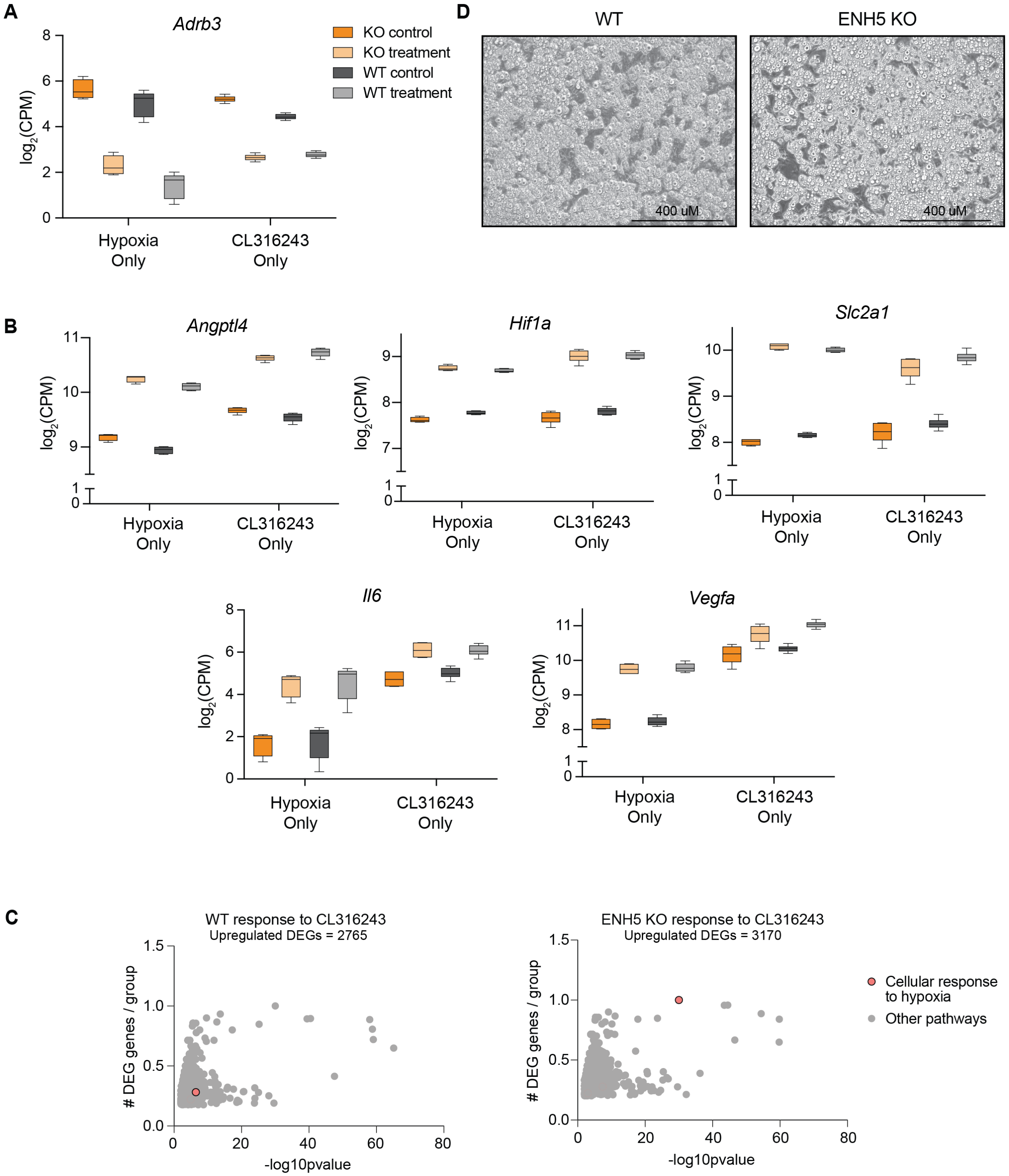
